# Identification of Common Adulterants in Walnut Beverage Based on Plant DNA Barcode Technology

**DOI:** 10.1101/340612

**Authors:** Qing Han, Yan Zhang, Ying Shu, Jia Chen, Yalun Zhang, Wei Zhou, Zhisheng Zhang

## Abstract

Walnut beverage is a common vegetable protein drink that is rich in proteins and has a wide consumption market. In this study, a plant DNA barcode technology was used to establish a method to identify common adulterated ingredients (peanut and soybean) in walnut beverage. In this experiment, universal primers were designed, and PCR amplification was performed. The universal primers were screened by sequencing and comparing the amplified products. Results showed that the primers rbcL-4 and matK-4 amplified walnut, peanut, sesame, soybean, and hazelnut. Peanut genomic DNA and soybean genomic DNA were added to the genomic DNA of walnut in different proportions. Primer rbcL-4 can detect 10% peanut genome DNA, and primer matK-4 can detect 10% soybean genome DNA. Calculation results of the extraction rate revealed that primer rbcL-4 can detect 8.88% peanut raw materials and primer matK-4 can detect 2.30% soybean raw materials. The combination of these two primers can be used as a universal primer for the identification of adulterated components in walnut beverage. This experiment could serve as a theoretical reference for related research and detection.

## Introduction

Walnut, also known Qiangtao, is rich in nutrients and is a favorite nut by consumers. Walnut is the main raw material for walnut beverage. After grinding, pulping, and other processes, water, sugar, and other prepared ingredients are added in the resulting slurry and then mixed into a milky beverage. Aside from pure walnut beverage, some compound vegetable protein beverages containing walnuts and milk ingredients are available in the market. As people’s living standards improve, the demand for these products is also increasing. However, some unscrupulous merchants deceive consumers through profit-driven adulteration and fraud. Therefore, methods to detect adulterants in these products are important.

DNA barcodes are standard, well-mutated, easily amplified, and relatively short DNA fragments that can represent the species in living organisms ^[1,2]^. DNA barcoding is a new technology for biological species identification ^[3–7]^. This technology has the advantages of high accuracy, wide detection range, and easy operation. Moreover, sequence stability is not changed by individual development. This technology could be effective for species identification in the future. At present, COI genes such as *ITS, matK, rbcL,* and *trnH-psbA* are mostly used in the identification of animal species ^[8–10]^.

Some studies have focused on detecting adulterants of walnut beverage. Wei Xiaolu ^[11]^ established a rapid detection method for peanut and soybean in walnut beverage by designing specific primers for walnut, peanut, and soybean. Zhang Jingjing ^[12]^ used gas chromatography and high-performance liquid chromatography to detect behenic acid from peanut and catherine genistein from soybean in walnut beverage, respectively. Yang Shuo ^[13]^ established a detection method for walnut and soybean components in walnut beverage by using multiple digital PCR. This study established a strategy based on DNA barcoding to detect common adulterants (peanuts and soybeans) in walnut beverage. This study could serve as a theoretical reference for relevant research and testing departments.

## Materials and methods

### Materials and reagents

Almonds, peanuts, walnuts, soybeans, sesame, hazelnut, and Yiyuantian, Xueming, Lvling, Yangyuan, Yili, Leliuren, Sijiyangguang, Aerfa, Chunwang, Zhihuiren, Moernongzhuang, Hezhilin, Yangbi, Mojiang, Fumanjia, Aibaren, Rikang, Jiajin, and Haobangxiang walnut beverages were purchased from Shijiazhuang Free Market of Agricultural Products and Supermarket.

Deep-processed Food DNA Extraction Kit was purchased from Tiangen Biotech Co., Ltd. (Beijing). Phenol:chloroform:isoamyl alcohol with a ratio of 25:24:1) was obtained from Coolaber Science and Technology Co., Ltd. (Beijing). In addition, isopropyl alcohol (analytical reagent), anhydrous ethanol (analytical reagent), and Premix TaqTM were purchased from Tianjin Guangfu Science and Technology Development Co., Ltd., Tianjin Oubokai Chemical Co., Ltd., and TaKaRa, respectively. 2xSuperReal PreMix (probe) was purchased Tiangen Biotech Co., Ltd. (Beijing). Meanwhile, specific primers and probes, universal primer, PCR product sequencing, and 50× TAE buffer were purchased from Sangon Biotech (Shanghai) Co., Ltd. GelRed™, Agarose, and 100 bp DNA ladder were purchased from Biotium, Vivantis, and Thermo Fisher Scientific Co,. Ltd, respectively.

### Instruments and equipments

An electronic balance (ME204/02), a desktop centrifuge, and a water bath constant temperature oscillator (SHA-B) were purchased from Mettler-Toledo Instrument Co., Ltd. (Shanghai), Sigma Centrifuge Co., Ltd. (Yangzhou), and Changzhou Runhua Co., Ltd, respectively. Moreover, a trace nucleic acid protein detector (NanoDrop Lite) and a fluorescence quantitative PCR system (LightCycler^®^ 480 II) were purchased from Thermo Fisher Scientific Co,. Ltd. and Roche Co., Ltd., respectively. A modular replaceable gradient PCR instrument (C1000 Touch™ Thermal Cycler), an electrophoresis system (PowerPac^TM^ Basic), and an almighty gel imaging system (ChemiDocTM MP Universal Hood III) were purchased from Bio-Rad Laboratories, Inc.

### DNA extraction

Sample genomic DNA was extracted using the optimized kit method. (1) Approximately 100 mg of the sample tissue was sufficiently ground with liquid nitrogen to sterilize the centrifuge tube. Afterward, 500 μL of buffer GMO1 and 20 μL of proteinase K (20 mg/μL) were added, and the solution was subjected to vortex shock for 1 min. (2) Then, the solution was incubated at 56 °C for 1 h, during which it was subjected to shock upside-down every 15 min. (3) Phenol, chloroform, and isoamyl alcohol were added at a ratio of 25:24:1. The resulting solution was mixed and centrifuged at 12,000 r/min for 5 min. (4) The upper water phase was transferred to a new centrifuge tube, and 200 μL of buffer GMO2 was added. Afterward, the solution was fully mixed, subjected to vortex shock for 1 min, and allowed to stabilize for 10 min. (5) The solution was centrifuged at 12,000 r/min for 5 min, and the upper water phase was transferred to a new centrifuge tube. (6) A total of 0. 7-fold higher volume than that of isopropanol was added into the supernatant. Then, it was fully mixed, cooled at −20 °C for 30 min, and centrifuged at 12,000 r/min for 3 min. Afterward, the supernatant was discarded, and the sediment was kept. (7) A total of 700 μL of 70% ethanol was added to the solution, and it was subjected to vortex shock for 5 s and centrifugation at 12,000 r/min for 2 min. Then, the supernatant was discarded. (8) Step 7 was repeated. (9) The lid was opened to dry the residual ethanol at room temperature. (10) A total of 50 μL of elution buffer TE was added, and the solution was subjected to vortex shock for 1 min.

The extraction method of the walnut beverage sample is as follows. (1) A total of 2 mL of the sample was placed in a centrifuge tube, and an equal volume of isopropanol was added. Afterward, the solution was mixed, allowed to stabilize for 15 min, and centrifuged at 12,000 r/min for 10 min. Then, the supernatant was discarded. (2) Step 1 was repeated. The rest of the steps are the same as the above-mentioned extraction process.

### Determination of sample accuracy

To ensure sample accuracy, NanoDrop Lite was used to determine the A260/A280 value and concentration of the extracted genomic DNA. The extracted genomic DNA was used as a template to amplify real-time fluorescence PCR with specific primers and probes of almonds, peanuts, walnuts, soybean, sesame, and hazelnut (SN/T 1961.9-2013, SN/T 1961.2-2007, SN/T 1961.6-2013, SN/T 1961.19-2013, SN/T 1961.12-2013, and SN/T 1961.8-2013, respectively). Moreover, amplification was performed to confirm the accuracy of the samples. PCR amplification of the genomic DNA was performed for a 25 μL volume reaction containing 12.5 μL of 2× SuperReal PreMix (probe), 0.75 μL of sense primer (10 μmol/L), 0.75 μL of antisense primer (10 μmol/L), 0.5 μL of probe (10 μmol/L), 2.0 μL of DNA template (30–100 μg/mL), and 8.5 μL of ddH_2_O. The reaction procedure was predenaturation at 95 °C for 15 min, and two major steps were repeated for 40 cycles, that is, (1) denaturation at 95 °C for 3 s and (2) annealing/extension at 60 °C for 20 s and collecting fluorescence signal.

### Primer design

Primers used were *ITS-1, ITS-2, rbcL-1, matK-1, matK-2, matK-3, and trnH-psbA-1*^[4–19]^. The primer sequences are shown in Table 1.

Primers *rbcL-2, rbcL-3, rbcL-4, matK-4, and matK-5.* The sequence of corresponding genes of each species was retrieved from NCBI. The software DNA MAN was used for comparison, and the primers were designed using software Primer Premier 5.0. The primer sequences are shown in Table 1.

**Table 1.**
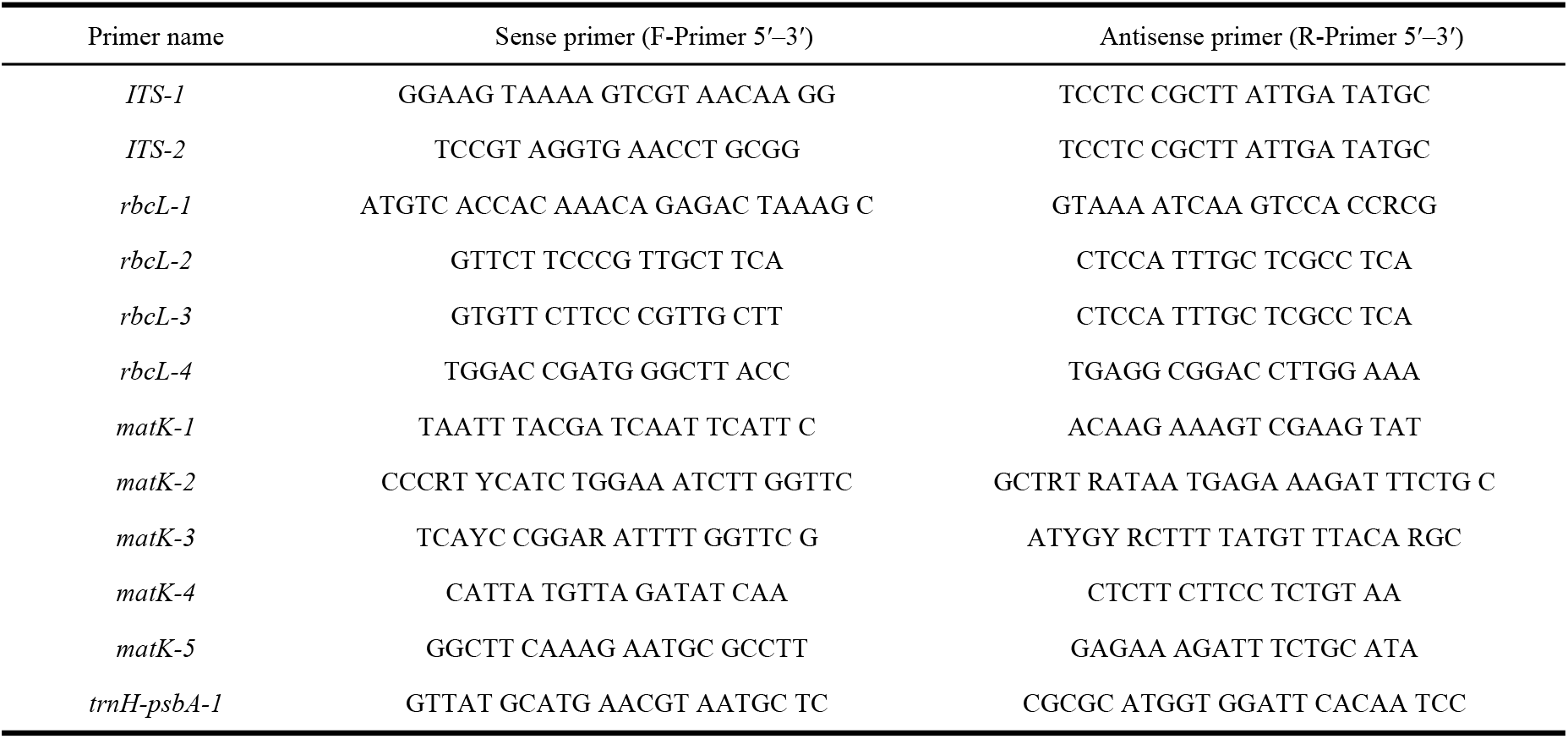
Primer sequences of genes

### PCR amplification

PCR amplification of the genomic DNA for a 50 μL volume reaction containing 25.0 μL of Premix Taq^TM^, 2.0 μL of sense primer (10 μmol/L), 2.0 μL of antisense primer (10 μmol/L), 4.0 μL of DNA template (30–100 μg/mL), and 17 μL of ddH_2_O was performed. The reaction procedure was as follows: predenatured at 95 °C for 5 min; 40 cycles of denaturation at 95 °C for 30 s, annealing at 45 °C (rbcL-2, rbcL-3, matK-4, and matK-5), 48 °C (ITS-1 and matK-3), 50 °C (rbcL-1, matK-1, and matK-2), 55 °C (ITS-2 and trnH-psbA-1), and 63 °C (rbcL-4) for 30 s, and extension at 72 °C for 40 s; and a final extension at 72 °C for 10 min. After the reaction was completed, agarose gel electrophoresis detection of 7 μL PCR amplification products was performed.

### Sequencing and comparison of PCR products

PCR amplification products were frozen at −20 °C and loaded on an ice pack into the incubator in advance to be sent to Sangon Biotech Co., Ltd. (Shanghai) for bidirectional sequencing. Sequencing primers are the same as the amplification primers. The sequenced results delete both primer sequences. Thus, BLAST alignment was performed on NCBI, and the sequence of the samples was identified and analyzed in combination with the BOLD database (http://www.boldsystems.org).

## Results and discussion

### Determination of sample accuracy

The A260/A280 value and concentration of the extracted genomic DNA were determined. The A260/A280 value was 1.8–2.1, and the concentration of the control was 30–100 μg/mL, which is beneficial to subsequent experiments^[19]^. To ensure the accuracy of the samples, genomic DNAs extracted from almonds, peanuts, walnuts, soybean, sesame. and hazelnuts were identified by real-time fluorescence PCR. The results are shown in Figures 1–6.

**Fig. 1.**
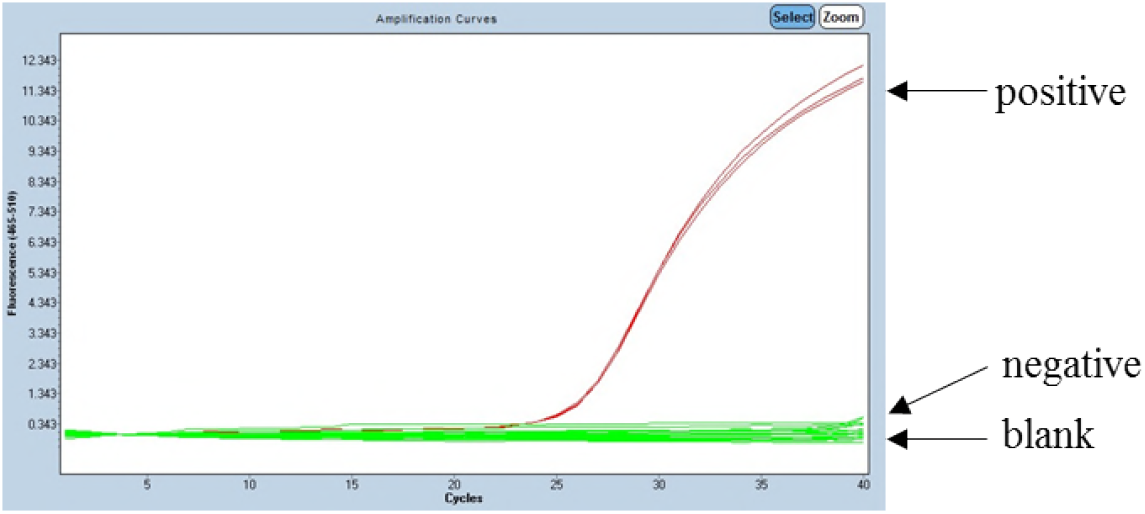
Real-time fluorescence PCR for the detection of almond genomic DNA

**Fig. 2.**
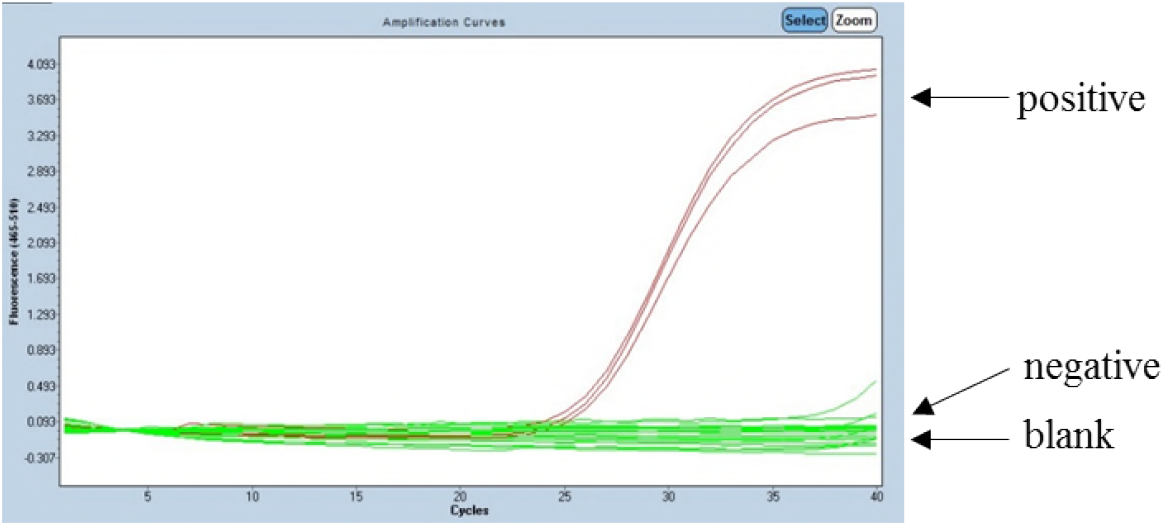
Real-time fluorescence PCR for the detection of peanut genomic DNA

**Fig. 3.**
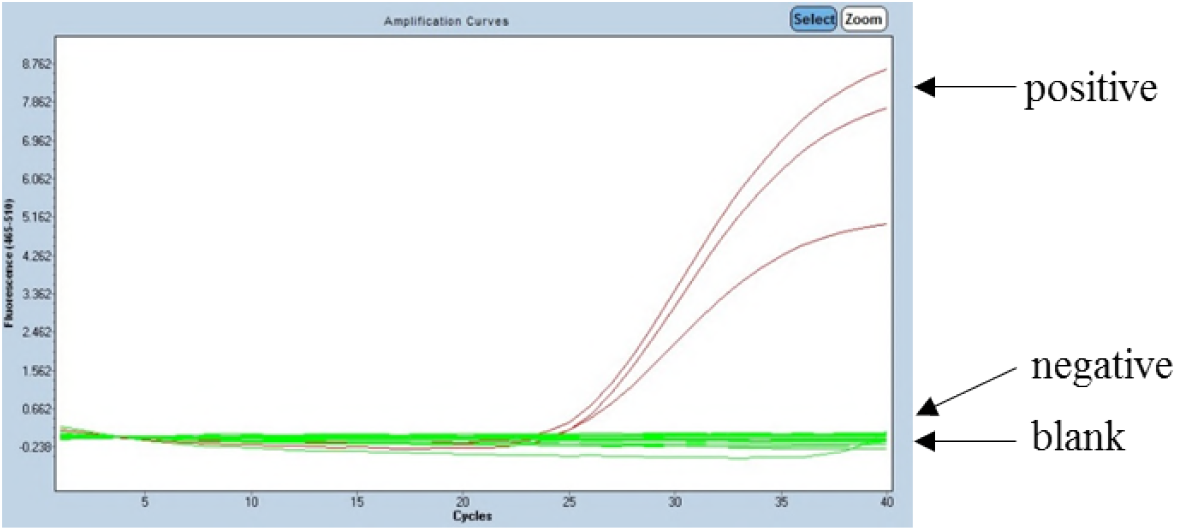
Real-time fluorescence PCR for the detection of walnut genomic DNA

**Fig. 4.**
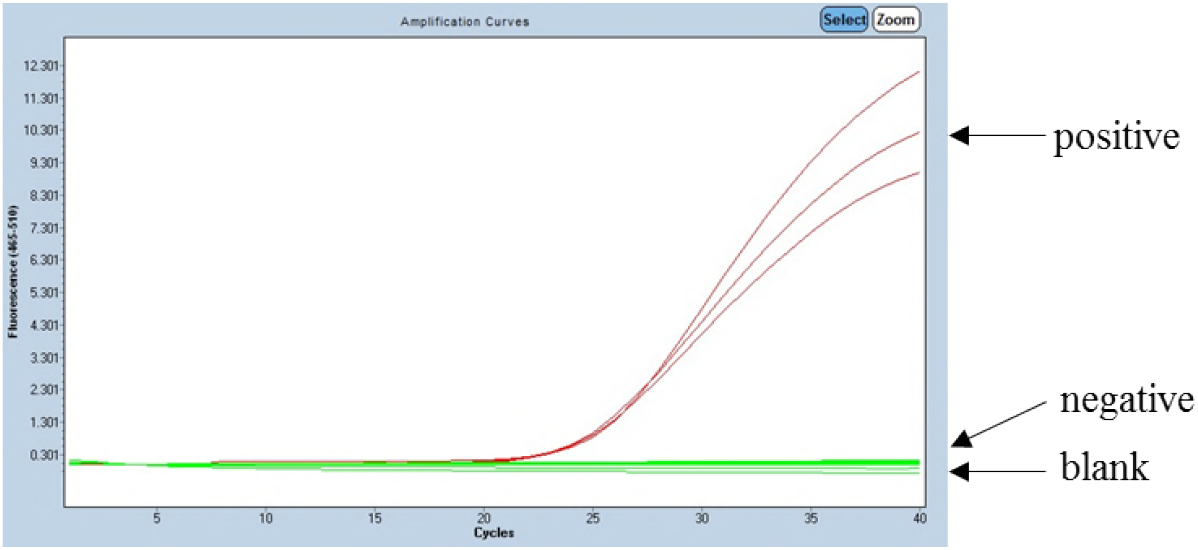
Real-time fluorescence PCR for the detection of soybean genomic DNA

**Fig. 5.**
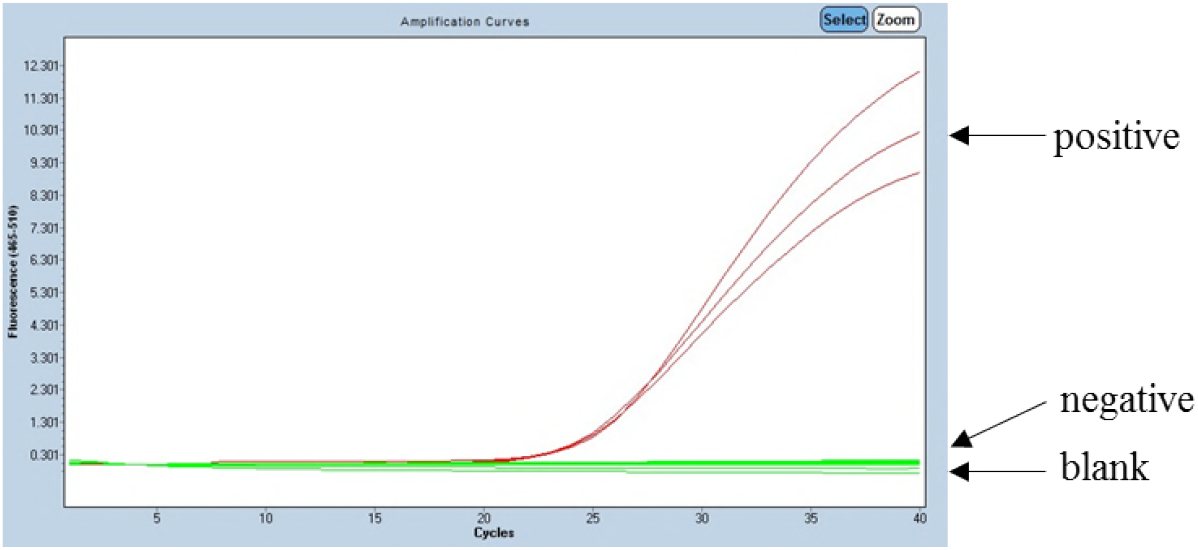
Real-time fluorescence PCR for the detection of sesame genomic DNA

**Fig. 6.**
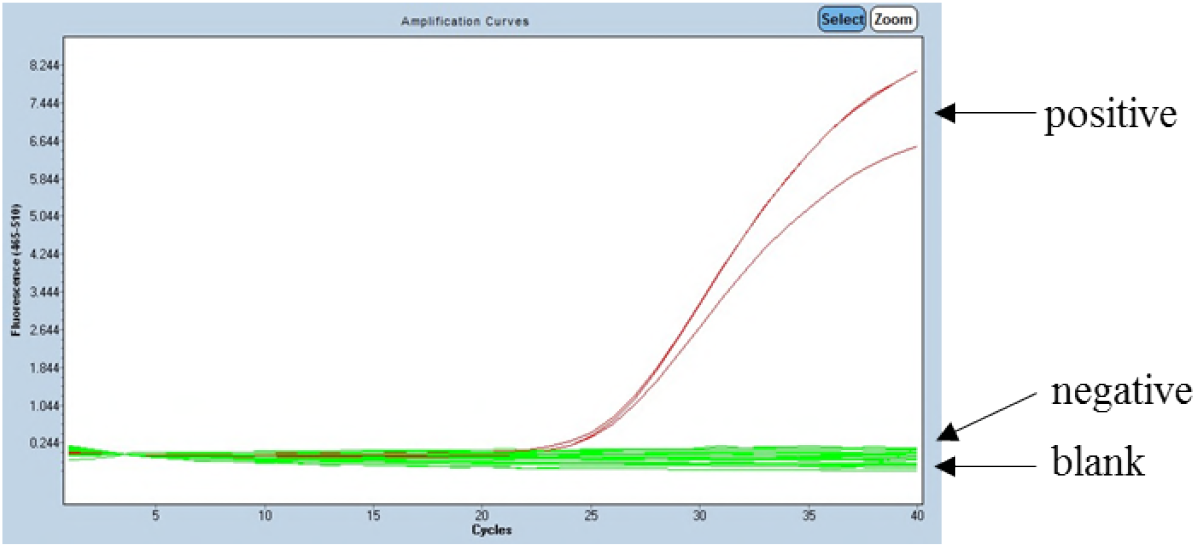
Real-time fluorescence PCR for the detection of hazelnut genomic DNA

With reference to the standard, Ct<30 (almonds and sesame), Ct≤35 (soybean, walnut, and hazelnut), and Ct ≤40 (peanut) were judged as positive. As shown in Figs. 1–6, the Ct values were all less than 30.0, and no negative or blank controls were detected. Therefore, all samples in this experiment were accurate and can be used in subsequent experiments.

### PCR amplification results

Primers were designed and referenced with six species of genomic DNA for PCR amplification. Sterile double distilled water was used as a blank control^[20]^. The amplified products were detected by agarose gel electrophoresis. After many optimization experiments, primers ITS-1, rbcL-2, rbcL-3, matK-1, matK-3, and matK-5 had low amplification efficiency for the six species used. These primers had poor universality and were excluded in subsequent experiments. Meanwhile, primers ITS-2, rbcL-1, rbcL-4, matK-2, matK-4, and trnH-psbA-1 can amplify the six species. The results of agarose gel electrophoresis are shown in Figs. 7–12.

**Fig. 7.**
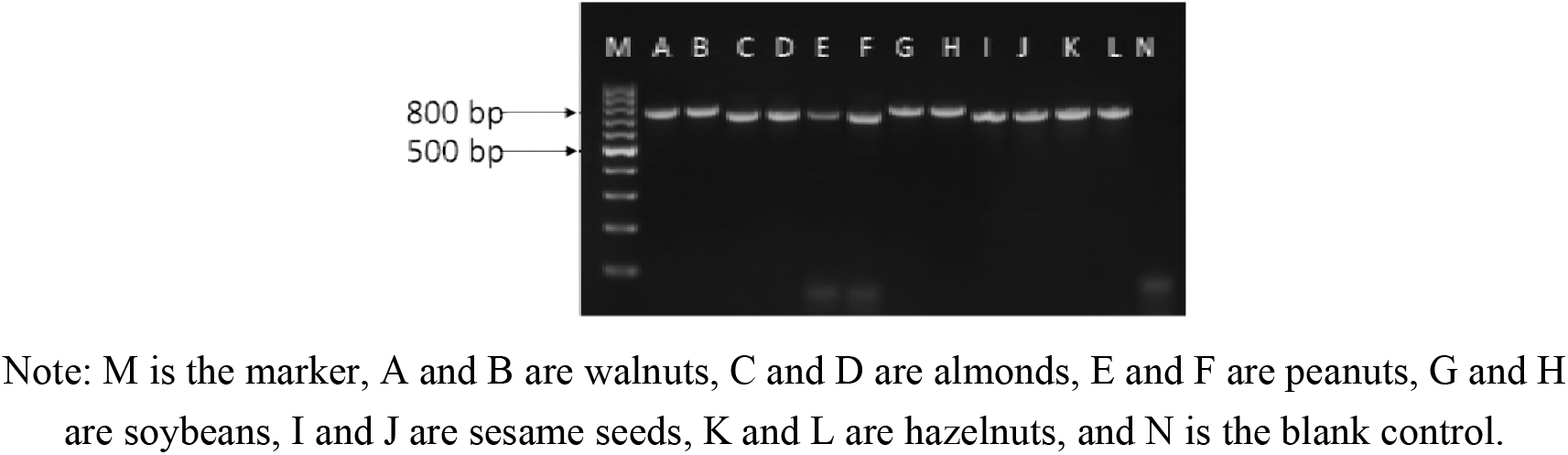
Specificity test result of primer ITS-2 in six samples

**Fig. 8.**
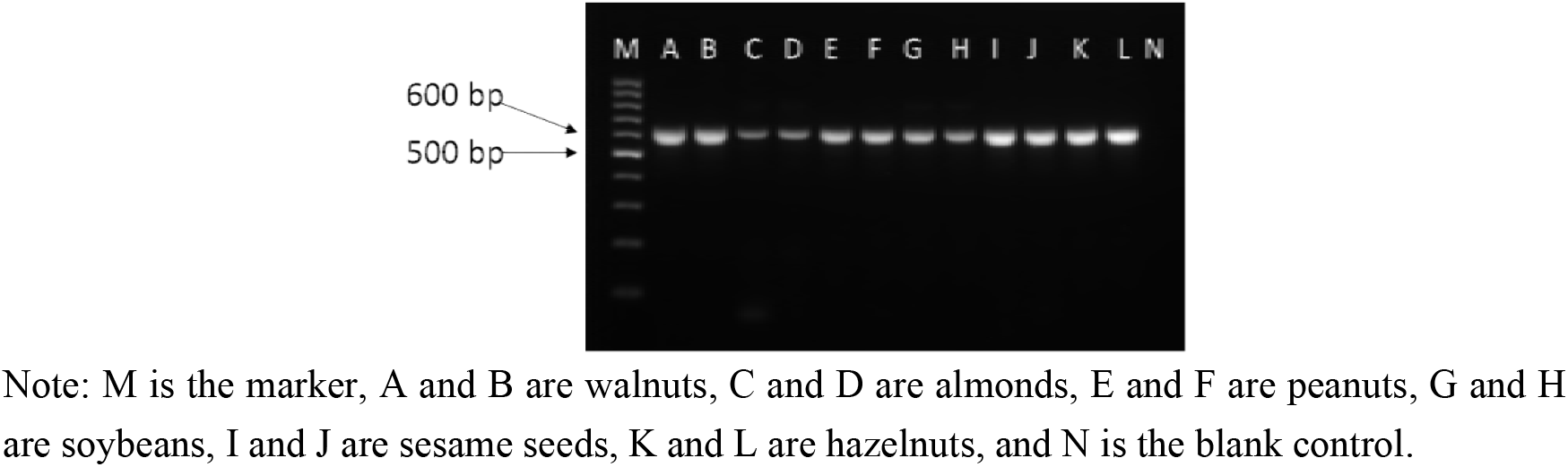
Specificity test result of primer rbcL-1 in six samples

**Fig. 9.**
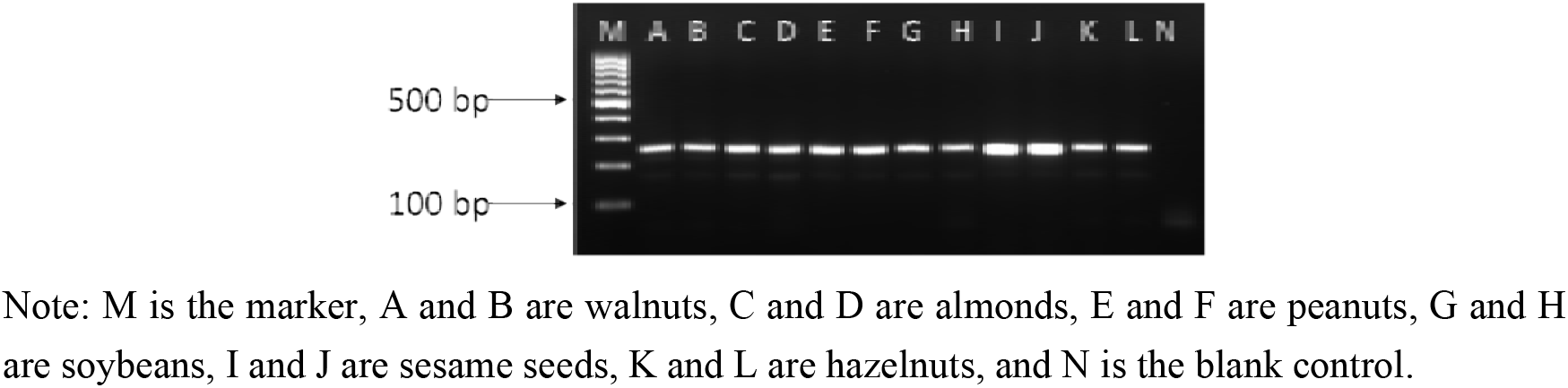
Specificity test result of primer rbcL-4 in six samples

**Fig. 10.**
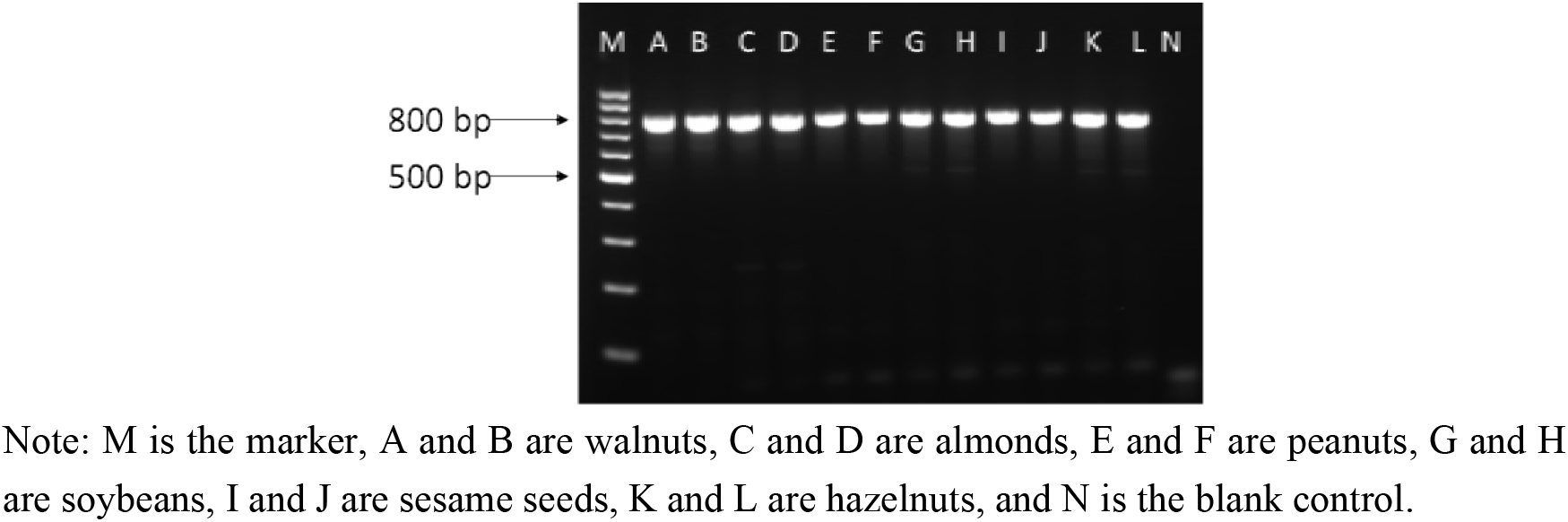
Specificity test result of primer matK-2 in six samples

**Fig. 11.**
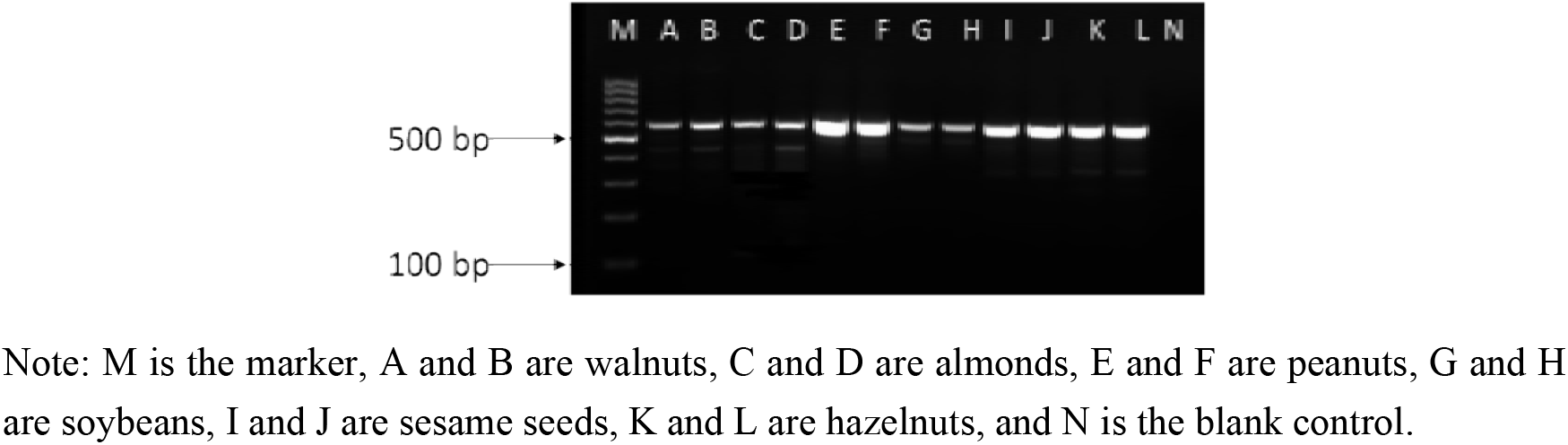
Specificity test result of primer matK-4 in six samples

**Fig. 12.**
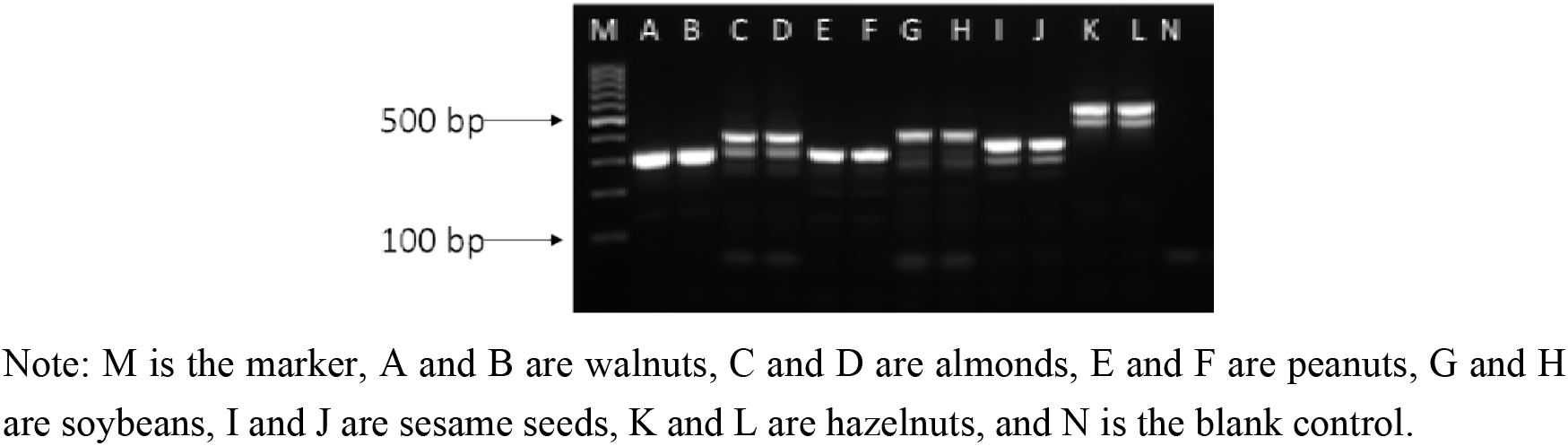
Specificity test result of primer trnH-psbA-1 in six samples

### Sequencing and analysis of PCR products

The PCR products were amplified with primers ITS-2, rbcL-1, rbcL-4, matK-2, matK-4, and trnH-psbA-1, and the six species were sent to Sangon Biotech Co., Ltd. (Shanghai) for bidirectional sequencing. The sequenced results deleted both primer sequences. BLAST alignment was performed on NCBI, and the sequence of the samples was analyzed in combination with the BOLD database^[21]^.

The results showed that primers rbcL-1, matK-2, and trnH-psbA-1 were successfully aligned, and their similarities were over 98%. Thus, these primers can be used as universal primers. Primer ITS-2 and the six species were all successful but were relatively low with the result of sesame comparison of 90%. Thus, ITS-2 can be used as the alternative universal primer for subsequent experiments. The results of primer rbcL-4 failed compared with those of almond and hazelnut, and the amplification results of primer matK-4 and almond and sesame failed as well. The results of their comparison with other species were all successful, and their similarities were all above 97%. For subsequent experiments mainly involving walnuts, peanuts, and soybeans, primers rbcL-4 and matK-4 can be used as alternative universal primers. Basing from the above-mentioned test process and result analysis, we conclude that primers ITS-2, rbcL-1, rbcL-4, matK-2, matK-4, and trnH-psbA-1 can be used for subsequent experiments.

### Establishment of adulterated model

The test cannot completely simulate the actual production. Thus, the sample used in the experiment is small. If adulteration is directly performed through the raw material, then it will cause significant error. DNA level adulteration was taken in this experiment, and then the raw material adulteration was obtained through conversion. Peanut and soybean genomic DNAs, which accounted for the total genomic DNA proportion of 90%, 80%, 70%, 80%, 50%, 40%, 30%, 20%, and 10%^[22]^, were added to walnut genomic DNA. The genomic DNAs of walnut, peanut, and soybean (125, 100, 75, 50, and 25 mg) were extracted. The extraction methods are described in Section 1.3.1, and the reagents with varying qualities were added proportionally. The extraction rate of the sample can be calculated from the DNA level to the raw material level. According to Equation 1, the extraction rate T of each sample was calculated as follows:

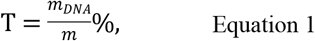

where T is the extraction rate, m_DNA_ is the total amount of DNA extracted from the sample, and m is the sample quality. When the genomic DNA was extracted from walnuts, peanuts, and soybeans of different qualities, three groups of parallel tests were performed. The extraction rates are shown in Table 2.

**Table 2.** Genomic DNA extraction rates of walnut, peanut, and soybean

| Sample name | Genomic DNA content<br>(ng) | Raw material<br>quality (mg) | Extraction rate<br>(%) | Average value<br>of extraction<br>rate (%) |
| --- | --- | --- | --- | --- |
| Walnut | 100895.83 | 125 | $8.07 \times 10^{-2}$ | $7.73 \times 10^{-2}$ |
| | 78840.00 | 100 | $7.88 \times 10^{-2}$ | |
| | 59210.00 | 75 | $7.90 \times 10^{-2}$ | |
| | 38311.67 | 50 | $7.66 \times 10^{-2}$ | |
| | 17852.50 | 25 | $7.14 \times 10^{-2}$ | |
| Peanut | 102066.67 | 125 | $8.17 \times 10^{-2}$ | $8.81 \times 10^{-2}$ |
| | 95806.67 | 100 | $9.58 \times 10^{-2}$ | |
| | 61811.25 | 75 | $8.24 \times 10^{-2}$ | |
| | 41746.67 | 50 | $8.35 \times 10^{-2}$ | |
| | 24245.83 | 25 | $9.70 \times 10^{-2}$ | |
| Soybean | 471025.00 | 125 | $37.68 \times 10^{-2}$ | $36.47 \times 10^{-2}$ |
| | 365160.00 | 100 | $36.52 \times 10^{-2}$ | |
| | 275280.00 | 75 | $36.70 \times 10^{-2}$ | |
| | 186472.50 | 50 | $37.30 \times 10^{-2}$ | |
| | 85315.00 | 25 | $34.13 \times 10^{-2}$ | |

According to Table 2, the genomic DNAs of walnut, peanut, and soybean were extracted using this method. The walnut extraction rate was 7.73 × 10^−4^, and the peanut extraction rate was 8.81 × 10^−4^. In addition, the extraction rate of soybean was 36.47 × 10^−4^. According to the extraction rates of walnut, peanut, and soybean and formula 1, conversion of DNA level to raw material level was carried out. In addition, the relationships between walnut and peanut and between walnut and soybean were obtained, as shown in Equations 2 and 3:

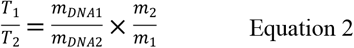

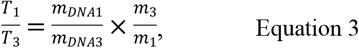

where T_1_ is the extraction rate of walnut, m_DNA1_ is the total amount of DNA extracted from walnut, and m_1_ is the quality of walnut. In addition, T_2_ is the extraction rate of peanut, m_DNA2_ is the total amount of DNA extracted from peanut, and m_2_ is the quality of peanut. Finally, T_3_ is the extraction rate of soybean, m_DNA3_ is the total amount of DNA extracted from soybean, and m_3_ is the quality of soybean. Taking each extraction rate into Equations 2 and 3, we obtained Equations 4 and 5 as follows:

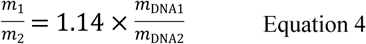

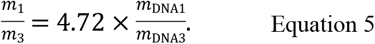

The DNA level of each adulteration ratio converted to the raw material level after the ratio is shown in Table 3 and 4.

**Table 3.** Conversion of DNA level to raw material level in walnut-mixed peanut model and the results of primer detection

| DNA level<br>adulteration<br>ratio (%) | Raw material<br>level adulteration<br>ratio (%) | Test<br>results<br>of ITS-2 | Test<br>results of<br>rbcL-1 | Test<br>results of<br>rbcL-4 | Test<br>results of<br>matK-2 | Test<br>results of<br>matK-4 | Test results<br>of trnH-psbA-1 |
| --- | --- | --- | --- | --- | --- | --- | --- |
| 10 | 8.88 | Walnut | Walnut | Peanut | Walnut | Walnut | Walnut |
| 20 | 17.99 | Walnut | Walnut | Peanut | Walnut | Walnut | Walnut |
| 30 | 27.32 | Walnut | Walnut | Peanut | Walnut | Walnut | Walnut |
| 40 | 36.90 | Walnut | Walnut | Peanut | Walnut | Walnut | Walnut |
| 50 | 46.73 | Walnut | Walnut | Peanut | Walnut | Walnut | Walnut |
| 60 | 56.82 | Walnut | Walnut | Peanut | Walnut | Walnut | Walnut |
| 70 | 67.18 | Walnut | Walnut | Peanut | Walnut | Walnut | Walnut |
| 80 | 77.82 | Walnut | Walnut | Peanut | Walnut | Walnut | Walnut |
| 90 | 88.76 | Walnut | Walnut | Peanut | Walnut | Walnut | Walnut |

Note: the proportion of the above-mentioned adulteration is the proportion of peanut in the whole.

**Table 4.** Conversion of DNA level to raw material level in walnut mixed soybean model and the results of primer detection

| DNA level<br>adulteration ratio<br>(%) | Raw material<br>level adulteration<br>ratio (%) | Test<br>results<br>of ITS-2 | Test<br>results of<br>rbcL-1 | Test<br>results of<br>rbcL-4 | Test<br>results of<br>matK-2 | Test<br>results of<br>matK-4 | Test results<br>of trnH-psbA-1 |
| --- | --- | --- | --- | --- | --- | --- | --- |
| 10 | 2.30 | Walnut | Walnut | Walnut | Walnut | Soybean | Walnut |
| 20 | 5.03 | Walnut | Walnut | Walnut | Walnut | Soybean | Walnut |
| 30 | 8.32 | Walnut | Walnut | Walnut | Walnut | Soybean | Walnut |
| 40 | 12.38 | Walnut | Walnut | Walnut | Walnut | Soybean | Walnut |
| 50 | 17.48 | Walnut | Walnut | Walnut | Walnut | Soybean | Walnut |
| 60 | 24.12 | Walnut | Walnut | Walnut | Walnut | Soybean | Walnut |
| 70 | 33.08 | Walnut | Walnut | Walnut | Walnut | Soybean | Walnut |
| 80 | 45.87 | Walnut | Walnut | Walnut | Walnut | Soybean | Walnut |
| 90 | 65.60 | Soybean | Walnut | Walnut | Walnut | Soybean | Walnut |

Note: the proportion of the above adulteration is the proportion of peanut in the whole.

The universal primer was screened with different levels of adulterated genomic DNA model for PCR amplification. Meanwhile, the amplified products were detected by agarose gel electrophoresis. Amplification of clear and bright bands of samples was sent to the sequencing company for sequencing.

As shown in Tables 3 and 4, the sequencing result was peanut when primer rbcL-4 was used to detect peanut genomic DNA (peanut material with 8.88% or above). Primer matK-4 was used to detect incorporation of 10% and above soybean genomic DNA (2.30% and the above-mentioned soybean material). Moreover, the sequencing results of other primers in different adulteration models were different for walnuts.

Test for screening out the lower detection limit and high sensitivity of universal primers for the detection of peanut and soybean, which are common adulterants in walnut beverage, was performed. Although primer rbcL-4 detects almonds and hazelnuts, and primer matK-4 detects almonds, sesame results are not ideal but can be applied to adulterated models compared with the other primers. At the early stage of experiment, we studied the adulteration of almond beverage by using plant DNA barcoding. Almond, hazelnut, and sesame were not commonly adulterated species in walnut beverage. At the same time, two pairs of universal primers can be used to detect hazelnut and sesame. When other primers were adulterated, the results of detection were still walnut when peanut or soybean with relatively high contents was added. When the primers were present, walnut was more competitive than the peanut and soybean primers, and the primers were easier to amplify with walnut. Thus, the combination of primers rbcL-4 and matK-4 was used as a universal primer.

## Sampling for testing and analysis

Genomic DNA was extracted from 30 samples of walnut beverage collected from different batches. The eukaryotic 18S rRNA gene was detected on extraction of walnut genomic DNA by real-time PCR amplification. The results are shown in Fig. 13. The figure shows that samples were takeoff, and the Ct value was <30.0, indicating that the sample genomic DNA conforms to the PCR amplification requirements.

**Fig. 13.**
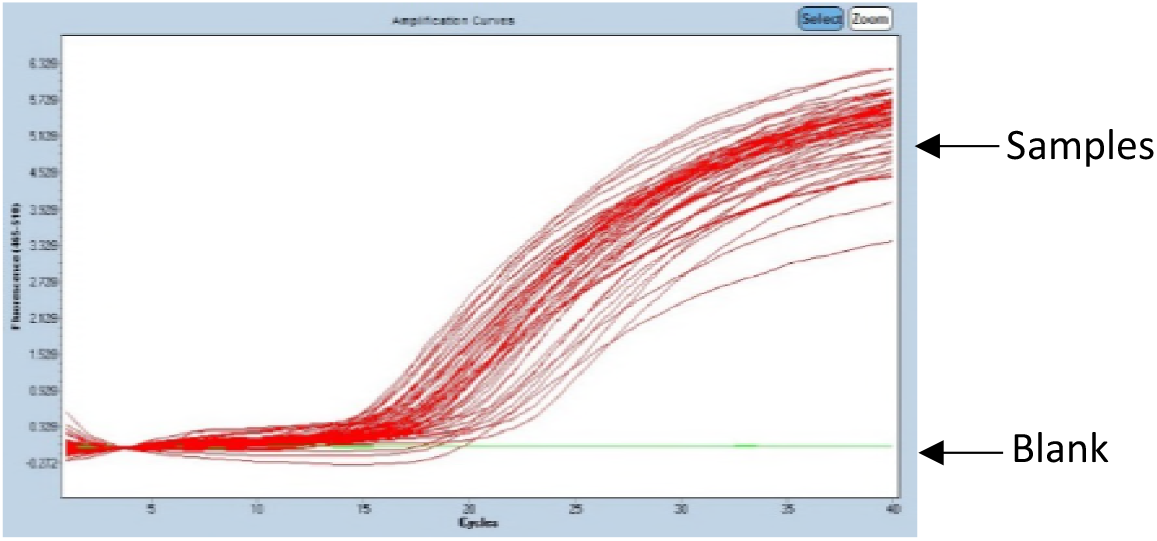
Real-time fluorescence PCR to detect the genomic DNA of walnut beverage

With primers rbcL-4 and matK-4, the extraction of walnut genomic DNA was amplified, wherein sterile double distilled water was used as blank control^[11]^. The PCR products were detected by agarose gel electrophoresis, and the results are shown in Fig. 14–17.

**Fig. 14.**
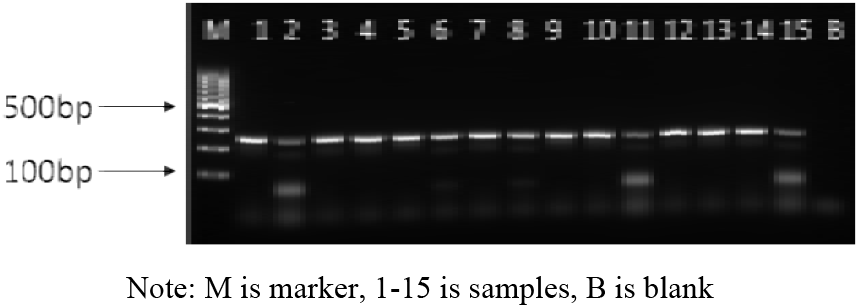
PCR result of primer *rbcL-4* in 1–15 walnut beverage

**Fig. 15.**
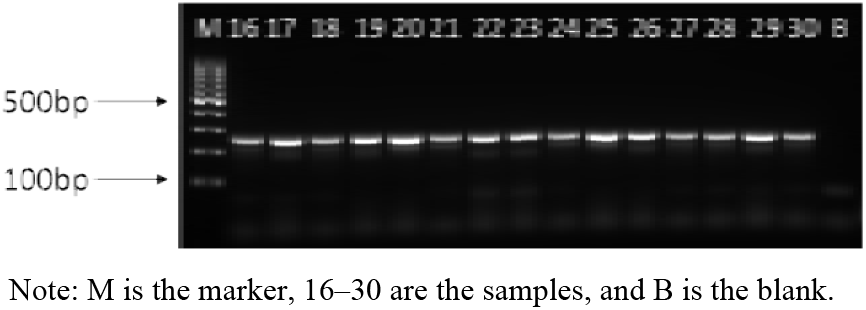
PCR result of primer *rbcL-4* in 16–30 walnut beverage

**Fig. 16.**
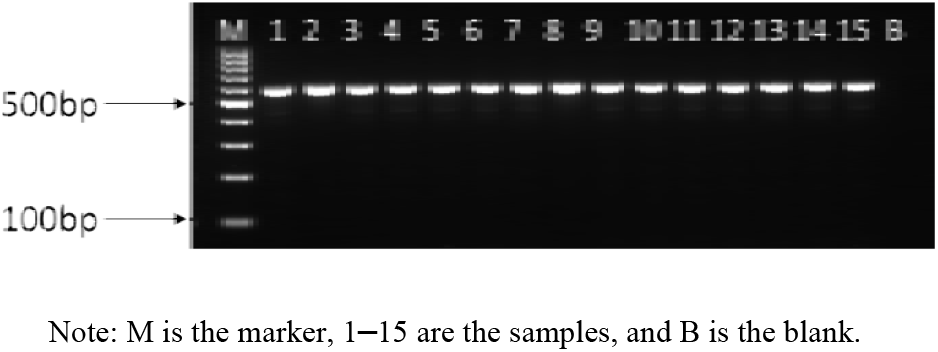
PCR result of primer *matK-4* in 1–15 walnut beverage

**Fig. 17.**
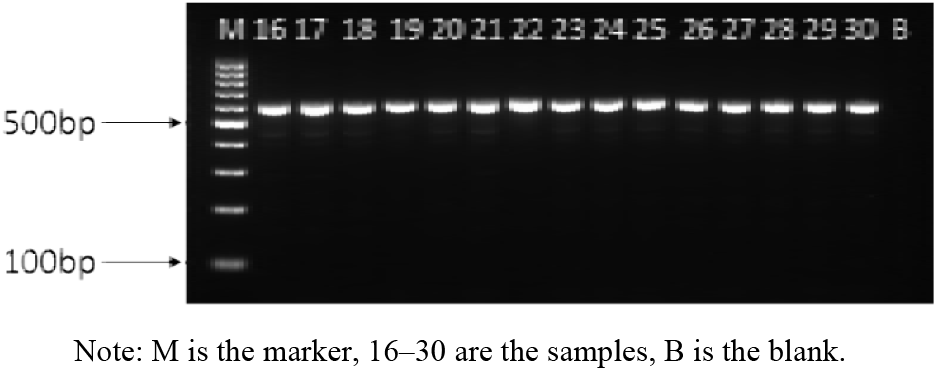
the PCR result of primer *matK-4* in 16-30 walnut beverage

As shown Figs. 14–17, 30 different batches of samples and 2 pairs of primers can be amplified by agarose gel electrophoresis results. The results showed that the bands were clear and bright, and no negative bands were observed. This result indicated that all PCR reaction system was polluted. The PCR amplification products of each sample were sent to Sangon Biotech Co., Ltd. (Shanghai) for bidirectional sequencing and sequencing result analysis. The sequencing results are shown in Table 5.

**Table 5.** Identification of samples

| Sample number | Plant source ingredients in the ingredient list | Comparison results of primer rbcL-4 detection | Comparison results of primer matK-4 detection | Identification results |
| --- | --- | --- | --- | --- |
| 1 | Walnut | Walnut | Walnut | Yes |
| 2 | Walnut | Peanut | Peanut | No |
| 3 | Walnut | Walnut | Walnut | Yes |
| 4 | Walnut | Walnut | Walnut | Yes |
| 5 | Walnut | Walnut | Walnut | Yes |
| 6 | Walnut | Peanut | Peanut | No |
| 7 | Walnut | Walnut | Walnut | Yes |
| 8 | Walnut | Peanut | Peanut | No |
| 9 | Walnut | Walnut | Walnut | Yes |
| 10 | Walnut | Walnut | Walnut | Yes |
| 11 | Walnut | Walnut | Walnut | Yes |
| 12 | Walnut | Walnut | Walnut | Yes |
| 13 | Walnut | Peanut | Walnut | No |
| 14 | Walnut | Walnut | Walnut | Yes |
| 15 | Walnut | Peanut | Walnut | No |
| 16 | Walnut | Walnut | Walnut | Yes |
| 17 | Walnut | Walnut | Walnut | Yes |
| 18 | Walnut | Walnut | Walnut | Yes |
| 19 | Walnut | Walnut | Soybean | No |
| 20 | Walnut | Walnut | Walnut | Yes |
| 21 | Walnut | Walnut | Walnut | Yes |
| 22 | Walnut | Peanut | Peanut | No |
| 23 | Walnut | Walnut | Walnut | Yes |
| 24 | Walnut | Walnut | Walnut | Yes |
| 25 | Walnut | Walnut | Walnut | Yes |
| 26 | Walnut | Walnut | Walnut | Yes |
| 27 | Walnut | Walnut | Walnut | Yes |
| 28 | Walnut | Walnut | Walnut | Yes |
| 29 | Walnut | Walnut | Walnut | Yes |
| 30 | Walnut | Walnut | Walnut | Yes |

According to the sequencing results of primers rbcL-4 and matK-4 shown in Table 5, samples 2, 6, 8, 13, 15, 19, and 22 did not match the labeled components of the product ingredient list. Moreover, peanut-derived ingredients or soybean-derived ingredients that were judged as adulterants after review of results of these seven batches were unchanged. The rest of the sample test results showed walnuts. Thus, no adulteration problem was present in the detection method.

## Conclusions

This experiment established a detection method for commonly adulterated ingredients in walnut beverage. Primers rbcL-4 and matK-4 had high efficiency and success rate of the amplification efficiency and sequencing of walnut, peanut, sesame, soybean, and hazelnut. The results of different proportions of adulteration model showed that primer rbcL-4 can detect 8.88% of peanuts mixed with walnuts, and matK-4 can detect 2.30% of soybean mixed with walnuts. In addition, the detected content was low. Using this method to test 30 batches of commercially available samples, we found that seven batches of samples had problems, including six spiked peanut samples and one spiked soy sample.

At present, relatively few reports on the adulteration of plant protein beverage are available. Some studies used species-specific primers for detection, and each species involved needs to be tested. Plant DNA barcoding uses universal primers for identification, with a relatively broader scope of application compared with other technologies. Some plant protein beverages include other ingredients, such as coconut, cashew nut, and pine nut. Moreover, primers rbcL-4 and matK-4 can amplify five species. Therefore, studying universal primers containing other species is important.

## Acknowledgments

This work was supported by Hebei Food Inspection and Research Institute. We thank Wang S, Zhang JJ, Li YH, Li YB for helpful guidance on an earlier experiment.

